# Injury Severity is a Key Contributor to Coagulation Dysregulation and Fibrinogen Consumption

**DOI:** 10.1101/2024.01.16.575945

**Authors:** Andrew R. Gosselin, Christopher G. Bargoud, Abhishek Sawalkar, Shane Mathew, Ashley Toussaint, Matthew Greenen, Susette M. Coyle, Marie Macor, Anandi Krishnan, Julie Goswami, Joseph S. Hanna, Valerie Tutwiler

## Abstract

**Background:** Traumatic injury is a leading cause of death for those under the age of 45, with 40% occurring due to hemorrhage. Severe tissue injury and hypoperfusion lead to marked changes in coagulation, thereby preventing formation of a stable blood clot and increasing hemorrhage associated mortality.

**Objectives:** We aimed to quantify changes in clot formation and mechanics occurring after traumatic injury and the relationship to coagulation kinetics, and fibrinolysis.

**Methods:** Plasma was isolated from injured patients upon arrival to the emergency department. Coagulation kinetics and mechanics of healthy donors and patient plasma were compared with rheological, turbidimetric and thrombin generation assays. ELISA’s were performed to determine tissue plasminogen activator (tPA) and D-dimer concentration, as fibrinolytic markers.

**Results:** Sixty-three patients were included in the study. The median injury severity score (ISS) was 17, median age was 37.5 years old, and mortality rate was 30%. Rheological, turbidimetric and thrombin generation assays indicated that trauma patients on average, and especially deceased patients, exhibited reduced clot stiffness, increased fibrinolysis and reduced thrombin generation compared to healthy donors. Fibrinogen concentration, clot stiffness, D-dimer and tPA all demonstrated significant direct correlation to increasing ISS. Machine learning algorithms identified and highlighted the importance of clinical factors on determining patient outcomes.

**Conclusions:** Viscoelastic and biochemical assays indicate significant contributors and predictors of mortality for improved patient treatment and therapeutic target detection.

**ESSENTIALS:**

- Traumatic injury may lead to alterations in a patient’s ability to form stable blood clots
- A study was performed to assess how trauma severity affects coagulation kinetics
- Key alterations were observed in trauma patients, who exhibit weaker and slower forming clots
- Paired with machine learning methods, the results indicate key aspects contributing to mortality

## INTRODUCTION

Traumatic injury is a leading cause of death both in the United States and globally, accounting for 180,000 and 5 million deaths respectively each year [1,2]. Among these traumatic injury deaths, 34-39% occur after arrival to the hospital [3,4]. Life threatening hemorrhage is estimated to account for 23%-39% of these trauma deaths and 40-90% of potentially survivable injuries [5-8]. Controlling life-threatening bleeding is dependent on timely physical intervention as well as the formation of a stable blood clot to mitigate blood loss at the site of injury. However, following severe traumatic injury this process is commonly impaired by Trauma Induced Coagulopathy (TIC), a multifactorial condition driven by the initial injury, development of hemorrhagic shock, and subsequent dysregulation of downstream factors [9-13]. TIC occurs in 25-35% of severely injured trauma patients resulting in a 4-6-fold increase in mortality and comprises both hypocoagulable and hypercoagulable phenotypes that lead to increased transfusion requirements and downstream thrombotic complications [9-13]. TIC is driven by the interplay of numerous factors including endothelial dysfunction, fibrinogen consumption, coagulation factor consumption, hypo- or hyperfibrinolysis, hemodilution from resuscitation fluids, platelet dysfunction, acidosis, and immune system dysregulation [10,12,13].

A stable blood clot is composed of platelets, red blood cells, white blood cells, other physiological components, and a polymeric fibrin network which provides the main mechanical and structural scaffold of the blood clot [14,15]. TIC leads to changes in the factors present in the blood, altering fibrin network formation, resulting in increases in maladaptive thrombosis or hemorrhage [9,14-16]. Alterations in the fibrin component of the blood clot and occurrence of fibrinolysis have been shown to impact clot structural mechanics and network properties [16-20]. Furthermore, changes to thrombin generation have been observed following injury in both animal models and trauma patients, leading to additional changes in fibrin structure and resulting clot stability [16,19,21,22]. TIC is known to contribute to mortality and altered clot formation, however the impact of traumatic injury on the coagulation system, TIC factors and clot stability have not yet been completely characterized [9,23].

Precisely characterizing how trauma induced coagulopathy occurs in injured patients is essential to direct life-saving interventions [24-26]. Viscoelastic and turbidimetric analysis allow for the real-time, qualitative assessment of coagulation kinetics, clot stability and presence of fibrinolysis [17,24,25]. A growing body of literature has demonstrated the value of viscoelastic testing at the bedside to inform treatment [24-26]. With the advent of machine learning and improved computational analysis methods, recent approaches have aimed to improve prediction and define which factors are most important in contributing to mortality following traumatic injury [27,28]. By pairing emerging methods of coagulation assessment with machine learning algorithms in this study, we aimed to characterize how injury severity contributes to coagulopathy progression, to define which mechanisms are associated with increased mortality, and which may be used for improved prediction.

## MATERIALS AND METHODS

### Patient Sample Collection and Preparation

Blood samples were collected from trauma patients (n=63) admitted to the Robert Wood Johnson University Hospital Emergency Department following informed consent under approval by the Rutgers’s University Institutional Review Board (Pro2021001296). All patients meeting the trauma center’s designation for the highest level of response necessary for care of life-threatening injuries were included in the study **(Table S1)**. After meeting inclusion criteria, whole blood (5 mL) was collected in 3.2% sodium citrate (9:1, v/v) from patients immediately after arrival to the resuscitation area. Blood was collected and stored at 4°C for a maximum of 16 hours before plasma was isolated. Control experiments were performed to confirm that collection and storage protocols did not alter the coagulability of samples. Platelet poor plasma was isolated by centrifugation at 2400 rpm for 10 minutes and stored at -80°C until subjected to mechanical, optical turbidity, ELISA, or confocal testing. After blood was collected and processed for storage, informed consent from the patient or surrogate was obtained as per Rutgers IRB approval. If inclusion criteria were not met or exclusion criteria were met due to over or under triage, or consent was not obtained, the sample was disposed of in accordance with institutional biosafety protocols.

### Healthy Individual Donor Sample Preparation

Healthy individual donors (n=7) had 15 mL of whole blood drawn following informed consent under the guidance of the Rutgers’s University Institutional Review Board (Pro2020001694) through venipuncture into 3.2% sodium citrate anticoagulant tubes (Grenier Bio, Kremsmunster, Austria). Platelet poor plasma was isolated by centrifugation at 2400 rpm for 10 minutes and frozen at -80°C prior to testing. For each testing method, samples were activated and supplemented in the same manner and at the same concentration as trauma samples.

### Rheometer Mechanical Testing

A Malvern Kinexus Ultra rheometer (Netzsch, Selb, Germany) was used to analyze the viscoelastic properties of plasma. Samples were activated with 20 mM CaCl_2_ and 0.6 Units/mL thrombin (final concentration), mixed, and transferred to the rheometer lower geometry. A 20 mm parallel plate oscillation strain test was performed as previously described [17].

### Optical Turbidity Testing

A Molecular Devices SpectraMax Plus (Molecular Devices, San Jose, California) plate reader was used to analyze the optical properties of plasma at 37°C. Turbidity measured the opacity of the solution as the plasma clot formed and lysed. Samples were activated as described above and tests were performed as previously described [17].

### Confocal Microscopy

A Zeiss LSM 710 Confocal microscope (Zeiss, Oberkochen, Germany) was used to image plasma from healthy and injured subjects. Clots were prepared in 500uL Eppendorf tubes, 1% of the plasma volume was replaced with 1.5mg/mL Alexa Fluor 488–labeled fibrinogen (Thermo Fisher Scientific, USA). Clotting was initiated as described above and samples were added to a clear bottom borosilicate glass 96 well plate and transferred to the confocal stage. Plasma samples were imaged using 1024×1024 resolution, 40x water immersion objective, 1.20 numerical aperture, with an image area of 213×213 micrometers. Image settings were kept constant across all images and samples. Z-stack images were obtained from three areas for each sample within 1 hour of clot activation with 11 slices imaged at an interval of 1 mm across a depth of 10 mm and maximum intensity projections were produced from these images.

### Image Analysis

Image analysis was completed using the Java ImageJ software version 1.8.0_322. The images were recorded as .czi files from the Zeiss Zen software and exported for ImageJ analysis. To calculate the area of fibrin density the files were made binary and total fiber area was measured as a percent of image area. Fiber length and pore diameter were measured manually using ImageJ and calculated from the average of 36 fibers and 16 pores for each sample.

### Determination of Plasma tPA Concentration

Tissue plasminogen activator (tPA) concentration in plasma was measured using the Abcam tPA human ELISA kit (ab108914, Abcam). Following the manufacturer’s instructions tPA was measured using a Molecular Devices Spectramax Plus (Molecular Devices, San Jose, California) plate reader, measuring absorbance at 450 nm wavelength and interpolation from the standard curve. Samples were diluted using provided diluent buffer, within the range of 1:8 dilution (1 part plasma, 8 parts diluent) to 1:2000 dilution (1 part plasma, 2000 parts diluent), to ensure chromogenic results were within standard range. All samples were run in duplicate.

### Determination of Plasma D-dimer Concentration

D-dimer concentration in human plasma was measured using the Abcam Human D-dimer ELISA kit (ab260076, Abcam). Following the manufacturer’s instructions D-dimer was measured using a Molecular Devices Spectramax Plus (Molecular Devices, San Jose, California) plate reader, measuring absorbance at 450 nm wavelength and interpolation from the standard curve. Samples were diluted using provided diluent buffer, within the range of 1:10 dilution (1 part plasma, 10 parts diluent) to 1:1200 dilution (1 part plasma, 1200 parts diluent), to ensure chromogenic results were within standard range. All samples were run in duplicate.

### Determination of Plasma Thrombin Generation

Thrombin generation was measured in patient and individual healthy donor plasma with the Technothrombin Thrombin Generation Assay (Diapharma, OH, USA) following the manufacturer instructions. The thrombin calibration curve was performed prior to plasma sample testing according to the manufacturer’s instructions. For all samples, 40 μL of sample was mixed with 50 μL TGA fluorogenic substrate and 10 μL TGA RC low trigger in duplicate. Fluorescence was immediately measured on a Molecular Devices SpectraMax M2 (Molecular Devices, San Jose, California) at 360 nm excitation and 460 nm emission wavelengths for 90 minutes at 1-minute intervals. The key parameters: lag time (LT), peak thrombin, time to peak (TTP), velocity index denoted in Relative Units (RU), and endogenous thrombin potential (ETP) were derived from the software provided from Diapharma.

### Determination of Plasma Fibrinogen Concentration

Plasma fibrinogen concentration was determined with the Dade Siemens Healthineers Fibrinogen Determination Kit (Siemens, Erlangen, Germany) following the manufacturer’s instructions using a STAGO Start 4 coagulation analyzer. Samples were warmed to 37 C, diluted 1:9 with Owren’s Veronal buffer (1 part plasma, 9 parts OV buffer) and incubated in the dilution cup for 2 minutes. Dade Thrombin Reagent (0.1 mL) was added after which clotting time was measured, with samples run in duplicate. Fibrinogen concentration was calculated based on the fibrinogen standard curve provided with the assay.

### Analysis using Machine Learning Methods

Binary classification using both clinical and biomechanical variables (n=103) was utilized to predict in-hospital trauma associated mortality. Methods applied included logistic regression with ridge regularization **(Table S2)** [29] (sklearn.linear_model library in python), support vector classifier [30,31] (sklearn.svm), feedforward neural network [32] (tensorflow.keras), and random forest [33] (sklearn.ensemble) chosen due to their relative edge [34] for classification problems. We tested 5 methods for comparative evaluation and identified the method that performed best on unseen test data. The data was split 60:40 between training and test cohorts and sampled randomly across the dataset with no order maintained when the split was done. A feedforward neural network was trained with four hidden layers and 100 epochs to reach convergence while training. The batch size of the input was set to 16. The optimization to reduce loss was done using Adam optimizer with a learning rate of 0.005. The dropout at each hidden layer was set to 50% to prevent overfitting and avoid training on noisy data. The activation function used on the hidden layers was the Rectified Linear Unit (ReLU) while the final layer had Sigmoid as the activation function. The architecture of the network was: Input: 103 x 128 x 64 x 32 x 8 x 1: Output. Logistic regression was employed using the sklearn.linear_model library in Python. Ridge regularization was used with a fit intercept and the L-BFGS solver. Support Vector Classifier was used with the regularization parameter set as 1.0, while the kernel used was linear and shrinking heuristic enabled (to speed up the training process). The training was done until the solver converged. The random forest algorithm which is an ensemble learning method (with a default of 100 trees) was applied. The grid search technique was employed to find the best hyper-parameters of the model. The hyper-tuned model had the following properties: the number of trees used were 200 where the maximum number of features considered at each level split was 20. The training samples were boot-strapped and out-of-bag samples were used as well to avoid overfitting and improving the generalization abilities of the model. Parallelization with 10 jobs was used for improved training speeds.

### Statistical Analysis

Statistical analyses were performed using GraphPad Prism 9.0. Test for normal distribution was assessed using a D’Agostino and Peason Normality test. One-way and two-way analysis of variance (ANOVA) were used to compare differences between data sets, followed by a Tukey test to determine statistical significance. Pearson correlation or nonparametric Spearman correlation analysis were performed, depending on normality, to determine correlations between groups. A nonparametric Spearman correlation matrix was used to compare each biomedical and key clinical factors, creating a heatmap of results. Simple linear regression analysis was performed to determine the proportion of variance of the impact of fibrinogen concentration on clinical and biomedical factors. All data are represented as the mean ± standard error of the mean unless otherwise noted for at least three replicates. Lack of significant differences between samples is indicated by no bar above the samples graphed. Rheological and turbidimetry clotting rate was defined as the rate of G’ or optical density increase from 10%-70% of the maximum sample value within 30 minutes of clot activation to reduce the impact of drying on analysis.

## RESULTS

### Patient Population Characteristics

Following Rutgers University Institutional Review Board (IRB) approval and informed consent, 63 patients between January 2021 and June 2023 were enrolled in the study, whole blood samples were drawn, and plasma was isolated as described. Patient characteristics, injury mechanism and pattern of all patients are shown in **Table 1**. The median age was 37.5 years old, 78% of the cohort was male, and the median Injury Severity Score (ISS) was 17 **(Table 1)**.

**Table 1:** Patient Injury Mechanism and Pattern Information.

|  | **Total (63)** | **Survived (44)** | **Deceased (19)** |
| --- | --- | --- | --- |
| **Median Age**<br>(Years) (25-75% IQR) | **37.5 (28.5-58)** | **33 (26-58)** | **47 (33-58)** |
| **Sex** (% male) | **49 (78%)** | **33 (75%)** | **16 (84%)** |
| **Injury Mechanism** |  |  |  |
| **Blunt** | **46 (73%)** | **29 (66%)** | **17 (89%)** |
| Motor Vehicle Collision | 22 | 16 | 6 |
| Fall | 11 | 7 | 4 |
| Pedestrian Struck | 10 | 6 | 4 |
| **Penetrating** | **17 (27%)** | **15 (34%)** | **2 (11%)** |
| Gun Shot Wound | 7 | 10 | 0 |
| Stab | 10 | 5 | 2 |
| **Median Injury Severity Score**<br>(ISS) (25-75% IQR) | **17 (10-29)** | **14 (9-26)** | **30 (17-52.5)** |
| **Injury Pattern** |  |  |  |
| Traumatic Brain Injury | 29 (46%) | 14 (32%) | 15 (79%) |
| Orthopedic injury | 4 (6%) | 4 (9%) | 0 (0%) |
| Spinal Cord Injury | 28 (44%) | 1 (3%) | 5 (26%) |
| Massive transfusion | 3 (5%) | 3 (7%) | 0 (0%) |

### Trauma Patients Exhibit Weaker and Faster Degrading Clots

Clot mechanical and functional fibrinogen testing showed that trauma patients exhibited significantly lower clot stiffness at 30 minutes (183.6 vs 80.1 Pa, p<0.0001) and fibrinogen concentrations (3.77 vs 2.54 mg/mL, p<0.01) compared to healthy donors **(Figure 1A,C)(Table S4)**. The injured had similar average clotting rate (0.23 vs 0.20 Pa/s) but larger variation in clotting rate compared to healthy donors **(Figure 1B)**.

**Figure 1:**
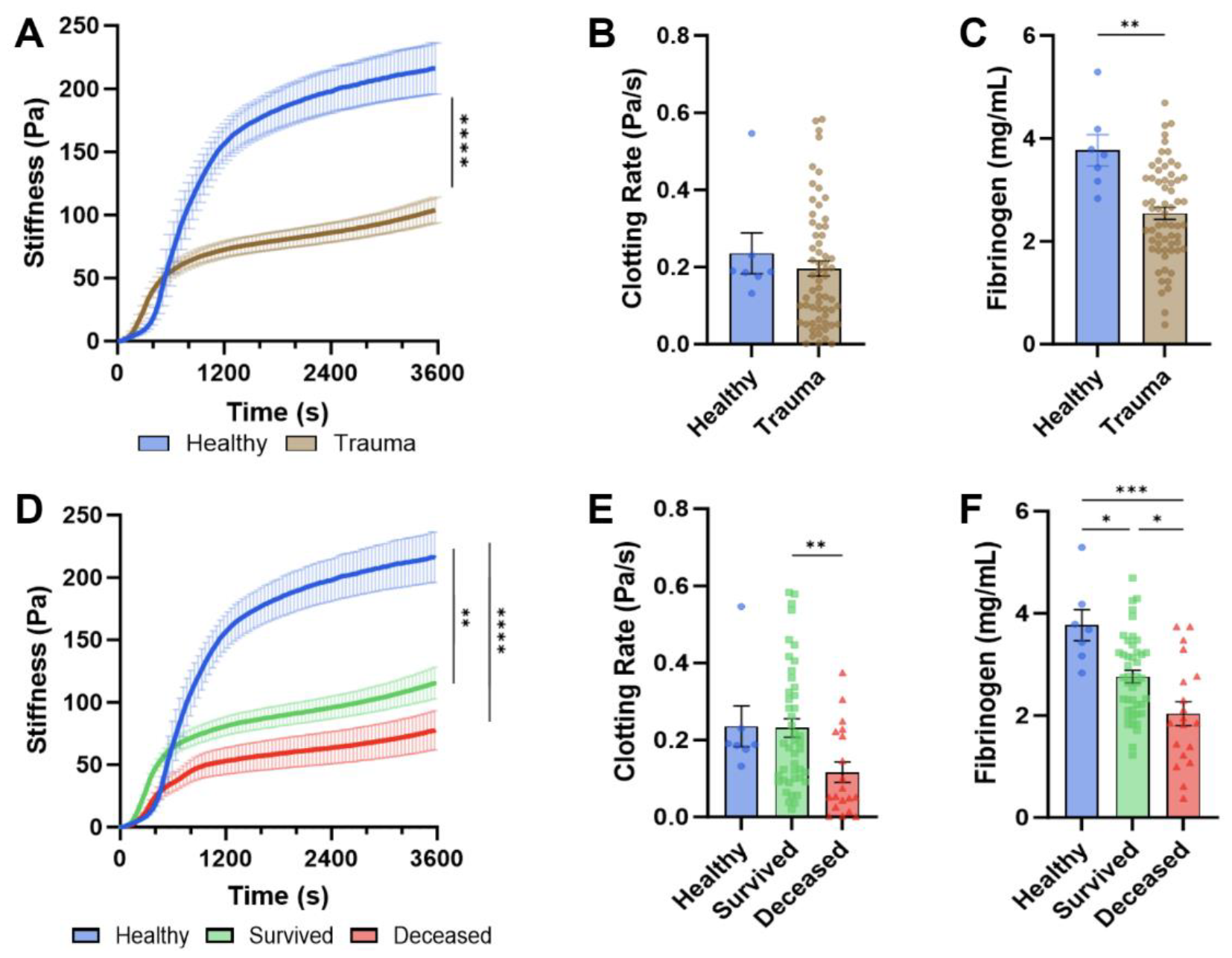
Trauma Patient and Healthy Donor Plasma Rheological Properties. Comparison of rheological parameters in healthy donors and trauma patients, stiffness over time **(A)**, clotting rate **(B)**, and fibrinogen concentration measured by clauss assay **(C**). Comparison between healthy donors, and patients (survived versus deceased), stiffness over time **(D)**, clotting rate **(E)** and fibrinogen concentration **(F)**. *Significance between groups indicated by* ^*\**^ *p < 0*.*05*, ** *< 0*.*01*, ^*\*\*\**^ *p < 0*.*001*, ^*\*\*\*\**^ *p < 0*.*0001*.

Turbidity testing revealed large differences in clot structure, with patients having reduced maximum optical density (0.82 vs 0.64 OD, p<0.01), and higher lysis after one hour (3.5 vs 12.1%, p<0.05), when compared to healthy donors **(Figure 2A,B)**. Trauma patients exhibited considerable upregulation of fibrinogen consumption prior to blood draw, with higher plasma D-dimer (0.48 vs 35.5 ug/mL, p<0.0001), compared to healthy donors **(Figure 2C)**. Paired with this, fibrinolysis following blood draw was similarly upregulated as shown with a significant mean increase in plasma tPA (2.4 vs 13 ng/mL, p<0.0001) **(Figure 2B,D)**. Further network changes were observed with confocal microscopy, though nonsignificant, an average lower fibrin fiber density, lower fiber length and larger pore diameter were observed in the trauma patient population, contributing to the optical density changes observed **(Figure S1A-C)**.

**Figure 2:**
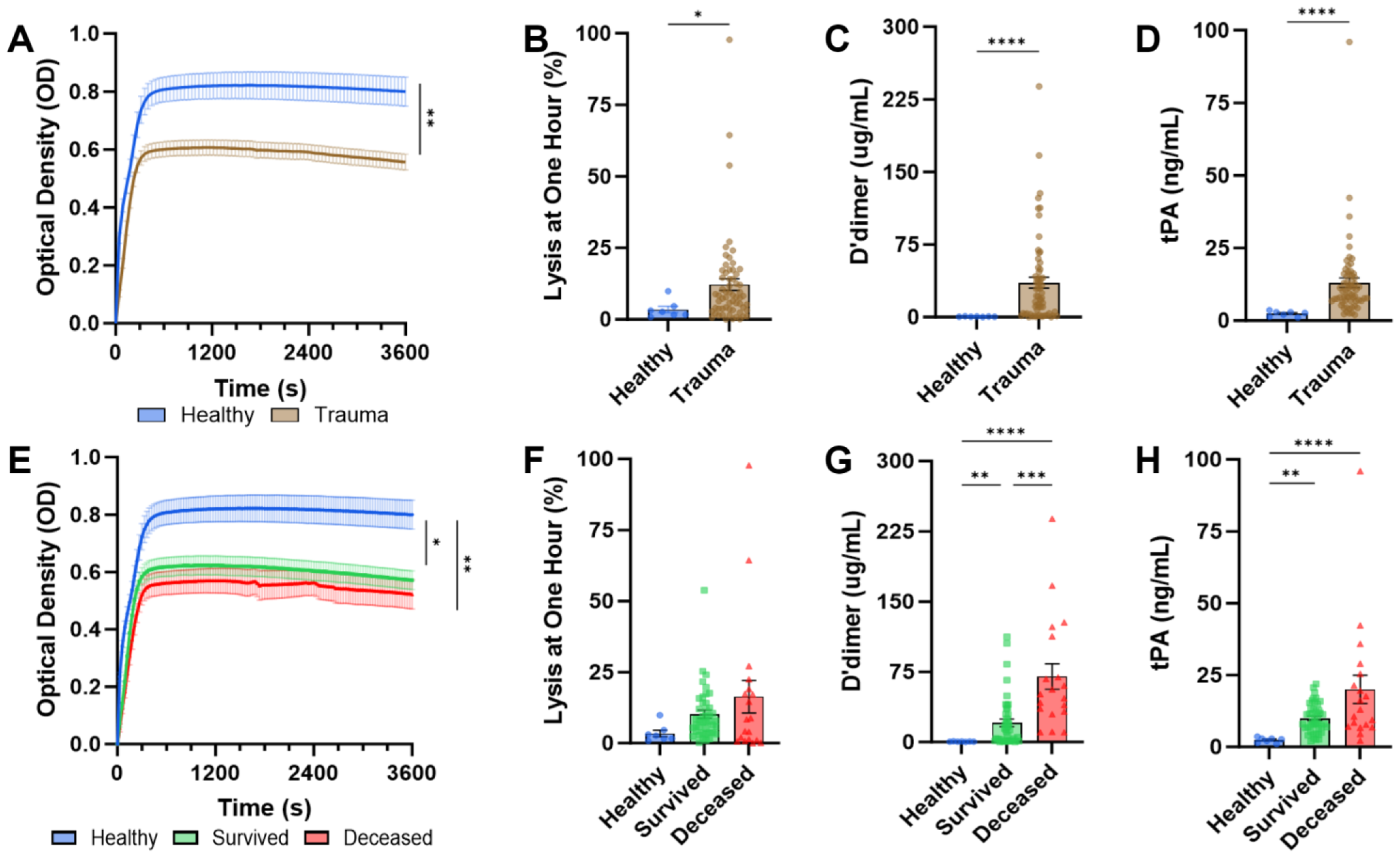
Trauma Patient and Healthy Donor Plasma Turbidity and Fibrinolysis Characteristics. Comparison between turbidimetric parameters in healthy donors and all trauma patients, measured by optical density over time **(A)**, lysis at one hour **(B)**, D-dimer concentration **(C)** and tPA concentration **(D)**. Comparison between healthy donors, and patients (survived and deceased), measured by optical density over time **(E)**, lysis at one hour **(F)**, D-dimer concentration **(G)** and tPA concentration **(H)**. *Significance between groups indicated by* ^*\**^ *p < 0*.*05*, ** *< 0*.*01*, ^*\*\*\**^ *p < 0*.*001*, ^*\*\*\*\**^ *p < 0*.*0001*.

Thrombin generation results indicate that on average, trauma patients exhibited increased lag time (8.4 vs 11 minutes, p<0.01), decreased peak thrombin (615 vs 488 nM, p<0.01), and decreased endogenous thrombin generation potential (ETP) (4752 vs 3891 nM, p<0.01) **(Figure 3A-D)**. Trauma patients also exhibited a prolonged time to peak thrombin concentration (TTP) (12.4 vs 15.4 minutes, p<0.01), while the rate of thrombin generation (Velocity Index) was only slightly reduced (163 vs 129 RU) **(Figure S1D**,**E)**. This data indicates significant thrombin generation attenuation seen in patients following severe injury.

**Figure 3:**
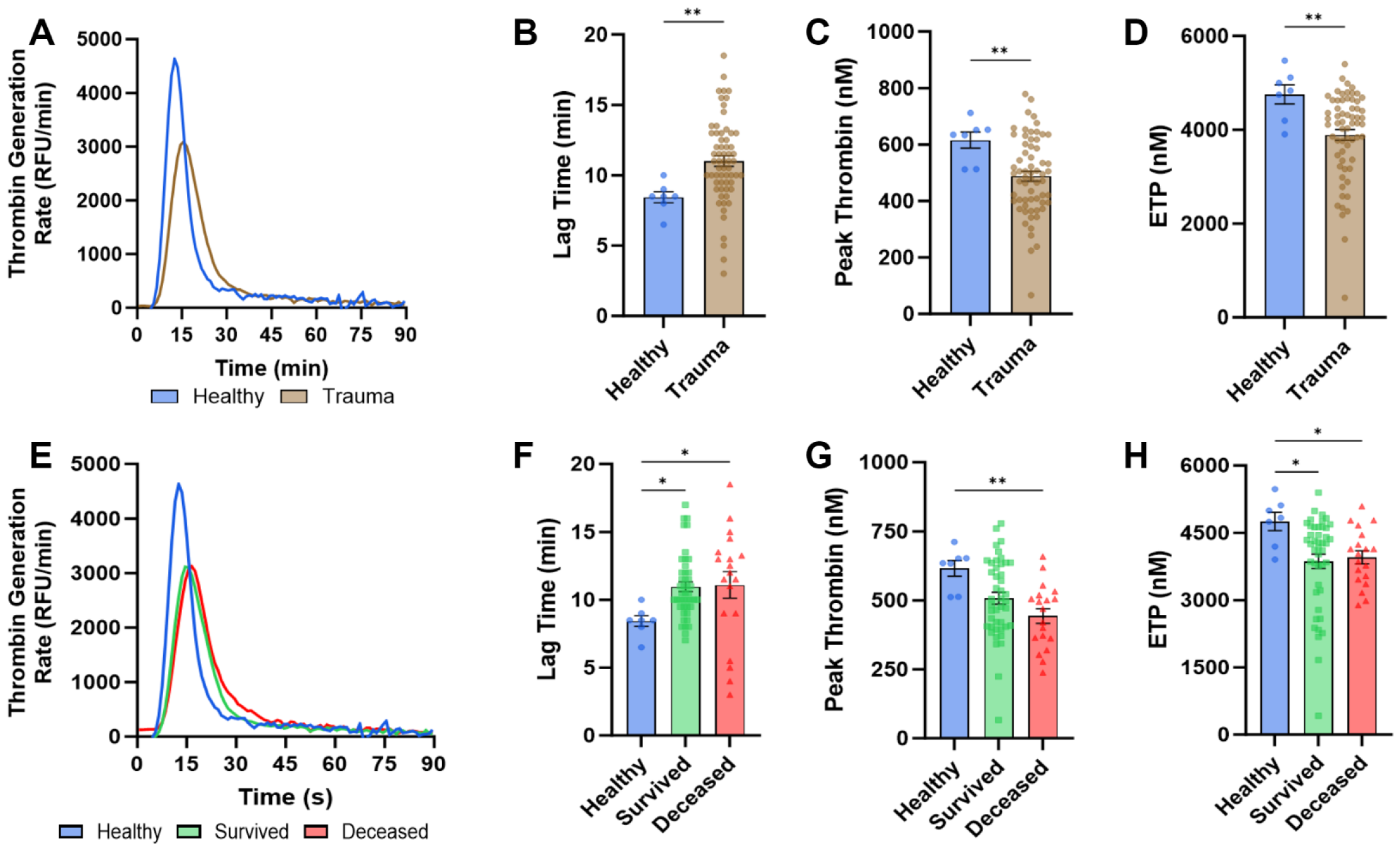
Trauma Patient and Healthy Donor Thrombin Generation Kinetics. Comparison between thrombin generation kinetics of healthy donors and all trauma patients, thrombin generation rate over time in relative fluorescent units per minute **(A)**, lag time to initial thrombin generation **(B)**, peak thrombin generated **(C)**, endogenous thrombin potential **(D)**. Comparison between healthy donors, and patients (survived and deceased), thrombin generation rate over time in relative fluorescent units per minute **(E)**, lag time to initial thrombin generation **(F)**, peak thrombin generated **(G)**, endogenous thrombin potential **(H)**. *Significance between groups indicated by* ^*\**^ *p < 0*.*05*, ** *< 0*.*01*.

### Deceased Patients Have Markedly Reduced Clotting Ability and Increased Factor Consumption

Significant differences were seen between healthy donors and trauma patients, as well as between patients who survived their injuries (survived) and those who died in hospital (deceased). Deceased patients were significantly more injured with higher Injury Severity Score (ISS) than patients who survived (p<0.0001) **(Figure 4B)**. Deceased patients exhibited reduced clot stiffness (89.1 vs 59.2 Pa) and significantly reduced clotting rate (0.23 vs 0.12 Pa/s, p<0.01) compared to surviving patients **(Figure 1D,E)**. Fibrinogen levels were also significantly lower in deceased patients (2.8 vs 2 mg/mL, p<0.01) than surviving patients and healthy donors **(Figure 1F)**.

**Figure 4:**
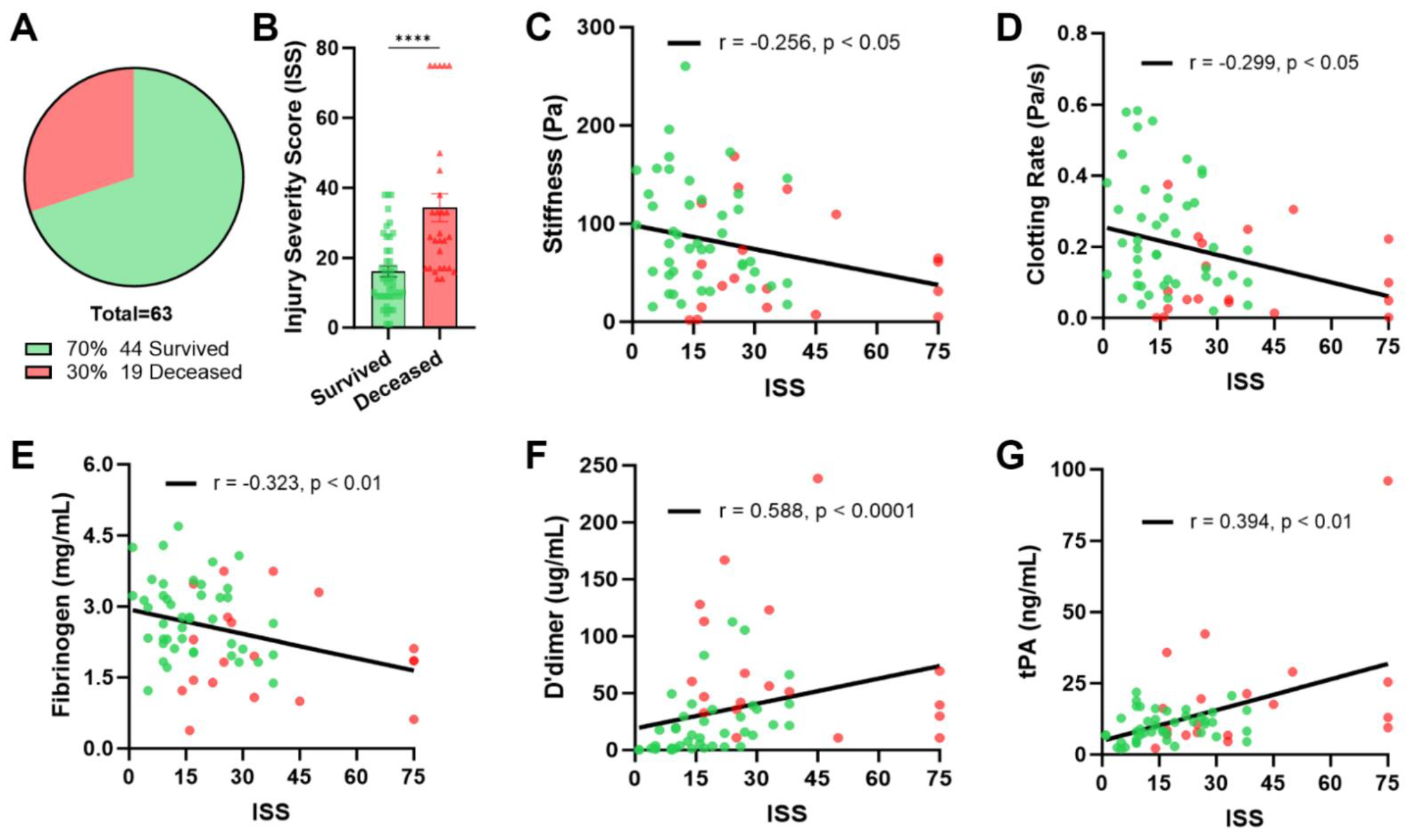
Correlation of Injury Severity Score with Coagulation Markers. Comparison of number of patients **(A)** and injury severity score **(B)** of survived and deceased patients. Correlation analysis of Injury Severity Score (ISS) and coagulation characteristics; maximum clot stiffness **(C)**, clotting rate **(D)**, fibrinogen concentration **(E)**, D’dimer **(F)** and tPA concentration **(G)**. *Significant correlations indicated by p < 0*.*05, r value indicates coefficient of correlation. Significance in t-test indicated by* ^*\*\*\*\**^ *p < 0*.*0001*.

Deceased patients had similar optical density but exhibited increased fibrinolysis at one hour (10.2 vs 16.4%), paired with higher tPA concentration (10 vs 20 ng/mL), compared to surviving patients **(Figure 2E,F,H)**. Deceased patients had increased lysis prior to blood draw, as indicated by increased D-dimer (20.6 vs 70.2 ug/mL, p<0.001) **(Figure 2H)**. Furthermore, deceased patients had the lowest average fibrin density and largest pore diameter observed by confocal microscopy **(Figure S1F-H)**.

Thrombin generation did not vary significantly between survived and deceased patients, with both having similar lag time (11 vs 11.1 minutes), peak thrombin (508 vs 443 nM), TTP (15 vs 16.6 minutes) and ETP (3864 vs 3954 nM) **(Figure 3E-H**,**S1I)**. The only significant difference present in thrombin generation was that deceased patients did have reduced velocity index compared to patients who survived (142 vs 98 RU, p<0.05) **(Figure S1J)**.

### Increasing Injury Severity Leads to Weaker Clots and Increased Consumption

With nearly one third of trauma patients dying from their injuries, and this group exhibiting a significantly higher mean injury severity score (p<0.0001), the correlation of ISS to coagulation kinetics and factor levels was assessed **(Figure 4A,B)**. Rheological analysis indicated a significant correlation of increasing ISS with a reduction in both maximum clot stiffness (p<0.05), clotting rate (p<0.05), and fibrinogen levels (p<0.05) **(Figure 4C-E)**. This loss of fibrinogen is correlated to the observed reduction in clot stiffness (p<0.0001) and clotting rate (p<0.0001) **(Figure 5B)**.

**Figure 5:**
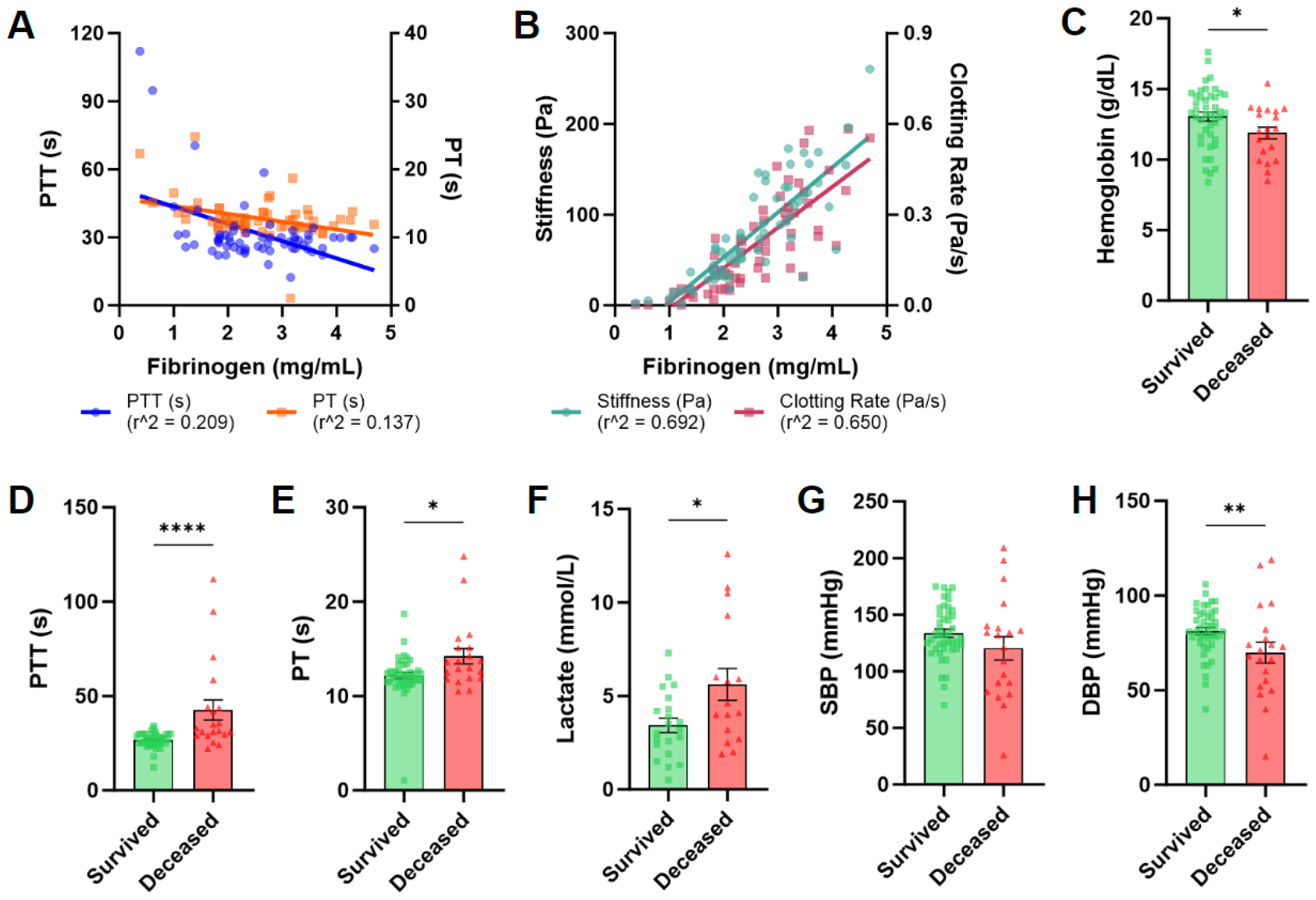
Patient Clinical Diagnostic Characteristics. Comparison of clinical parameters and fibrinogen concentration on clotting parameters. Association of fibrinogen concentration on clinical assays Partial Thromboplastin Time and Prothrombin Time **(A)** and impact of fibrinogen concentration on maximum clot stiffness and clotting rate **(B)**. Survived and deceased comparison of clinical parameters including Hemoglobin **(C)**, PTT **(D)**, PT **(E)**, Lactate **(F)**, Systolic blood pressure **(G)** and diastolic blood pressure **(H)**. *R^2 value indicates the coefficient of determination in linear regression analysis. Significance in t-test indicated by* ^*\**^ *p < 0*.*05*, ** *< 0*.*01*, ^*\*\*\**^ *p < 0*.*001*, ^*\*\*\*\**^ *p < 0*.*0001*.

Increasing ISS was not associated with significant changes in optical density or fibrinolysis at one hour **(Figure S2A**,**B)**. Though lysis did not correlate with ISS, fibrinogen consumption prior to blood draw, reflected in elevated D-dimer was correlated (p<0.0001) **(Figure 4F)**. Likewise, tPA release was positively correlated with increasing ISS (p<0.01) **(Figure 4G)**.

Confocal microscopy revealed that only fibrin fiber length significantly correlated to increasing ISS (p<0.05), whereas fibrin fiber density and pore diameter had only minor changes **(Figure S2C-E)**. Increasing ISS was not correlated with alterations in thrombin generation parameters **(Figure S2F-J)**. Lastly, to highlight the considerable interplay between factors, a correlation matrix analysis was performed on all measured coagulation parameters, as well as several clinical factors which reflect coagulation function **(Figure S3)**.

### Traditional Coagulation Assays Indicate Prolonged Coagulation in Deceased Patients

Trauma patients are routinely assessed for laboratory signs of coagulopathy to inform care and necessary surgical interventions. In the present analysis, prothrombin time (PT) and partial thromboplastin time (PTT) had limited correlation with increasing injury severity, or changing fibrinogen concentration, with low correlation r coefficients and R^2 regression coefficients **(Figure 5A,S3)**. In contrast, viscoelastic assays showed both high correlation to ISS and fibrinogen concentration, as well as high linear regression R^2 coefficients **(Figure 5B**,**S3)**. Deceased patients had a lower mean lower hemoglobin (p<0.05), lower diastolic blood pressure (p<0.01), and increased PTT (p<0.0001), PT (p<0.05), and lactate (p<0.05) **(Figure 5C-F,H)**.

### Machine Learning Identify Improved Methods to Predict Mortality

A total of 103 parameters were measured in the trauma population, including 77 clinical measurements in-hospital and 26 biomedical and other associated measurements in the laboratory. Five machine learning algorithms were used to identify features most predictive of in-hospital mortality. Accuracies of prediction were of the following: feedforward neural network 79.62%, baseline support vector classifier slightly higher at 81%, followed by baseline logistic regression and random forest, both at 85% accuracy, and the highest prediction noted in our hyper-tuned random forest model at 88% accuracy **(Table S3)**. The kernel density plots of top 4 of the 103 predictors of in-hospital mortality are shown in **Figure 6A**. The distinction in the mean and standard deviation between the classes of survived and deceased patients indicate their likelihood in making the predictions. The top 20 predictors of in-hospital mortality are shown in **Figure 6B**. The top 5 of which were D-dimer (19%), prothrombin time (PTT) (11%), rheological clotting rate (10%), fibrinogen concentration (9%) and head abbreviated injury score (6%) **(Figure 6B)**. Interestingly, 10 of the top 20 factors were laboratory measurements, contributing 57% of mortality prediction, driven by D-dimer and clotting rate **(Figure 6A-B)**. Both biomedical and clinical assays contribute unique aspects of mortality prediction with some biomedical factors being more predictive than current clinical standard practice.

**Figure 6:**
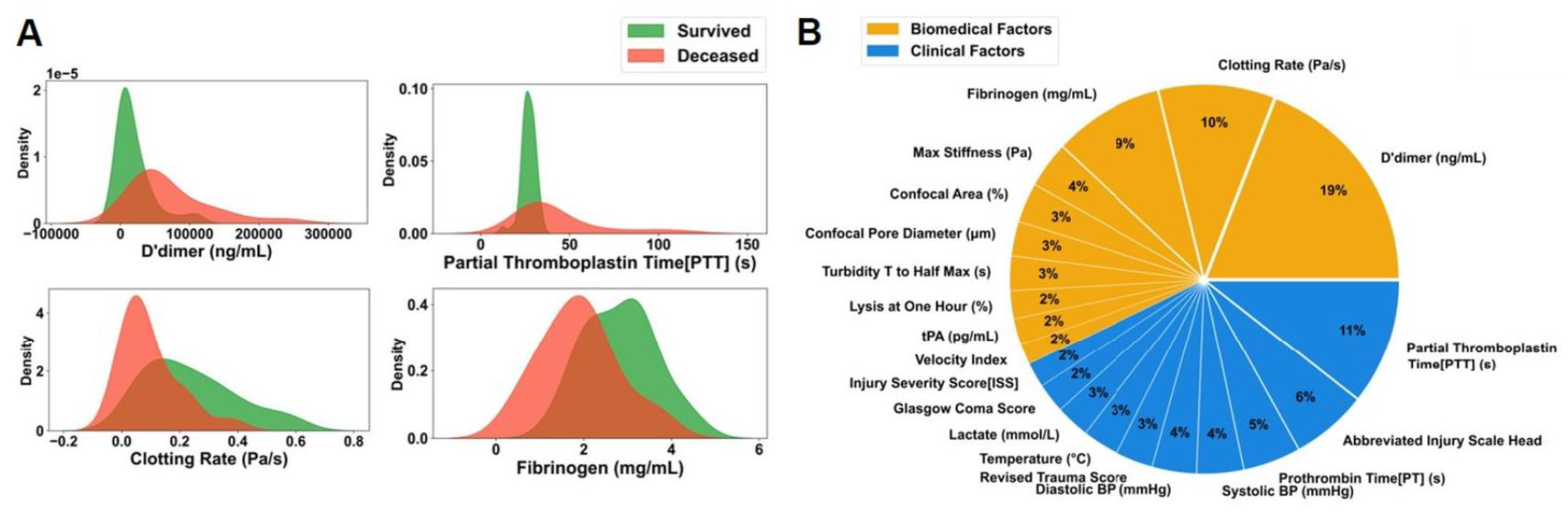
Key features associated with in-hospital mortality identified by random forest algorithms. **(A)** Kernel density plots of top 4 features significant in predicting patient mortality. Distinct mean and standard deviation of these features in patient mortality status survived vs deceased reflect model prediction accuracy for in-hospital mortality. **(B)** Pie chart representation of the top 20 of 103 total Biomedical and Clinical features contributing toward prediction of in-hospital mortality. Percent (%) contribution derived using Gini-index of the random forest model. *Biomedical values indicated in orange; clinical values indicated in blue*.

## DISCUSSION

This study aimed to characterize the correlation of injury severity and mortality with coagulation kinetics and clot mechanics to inform clinical interventions to reduce morbidity and mortality associated with trauma induced coagulopathy. By obtaining clinical samples prior to any intervention, our study allowed for the analysis of a snapshot of the coagulation milieu while minimizing the impact of iatrogenic confounders. The patient cohort studied had a wide variety of injury mechanisms, both blunt and penetrating, as well as injury patterns including soft tissue, brain, orthopedic and spinal cord injury **(Table 1)**. The study confirmed that injury severity plays a key role in promoting trauma induced coagulopathy. This was reflected in the observed decrease in fibrinogen concentration, upregulation of fibrinolytic markers, and reduction in thrombin generation **(Figure 1-3)**. These results, paired with novel machine learning techniques, expand our understanding of the mechanical perturbations to clot formation and emphasize key contributors to mortality which include D-dimer, PTT, viscoelastic clotting rate, fibrinogen, and head AIS. This analysis reinforces the value of real-time measurement to inform optimal bed-side clinical care.

### Severe Traumatic Injury Leads to Perturbations in Clot Stability

We identified an association between reduced clot stiffness and mortality, with significant correlation between increasing trauma severity, and fibrinogen loss resulting in reduced clot stiffness and clotting rate. Thus, not only is clot formation adversely affected, but any clots which do form may lack sufficient stability to support effective endogenous hemorrhage control. Thus, these data provide a plausible explanation for the observed mean decreased hemoglobin and diastolic blood pressure as well as increased lactate **(Figure 5C,F)** in the deceased patient cohort, and inform a more precise mechanistic understanding of the maladaptive physiologic changes occurring in the severely injured. Ultimately, the observed coagulopathy with resulting blood loss leads to insufficient oxygen delivery reflected in the observed lactic acidosis **(Figure 5C,F)**, two of the three physiologic disruptions associated with the triad of death [13].

Translating this to the bedside, without accounting for the increase in fibrinolytic activation and reduction in thrombin generation concomitantly seen in severely injured patients, treatment focusing solely on fibrinogen supplementation may be insufficient for rescue. However, we did not identify patient characteristics that reliably predicted the rate of fibrinolysis, limiting our ability to pre-emptively act to mitigate the adverse effects of TIC based solely on mechanism of injury **(Figure S2B**,**F-J)**. In response to injury, it is therefore essential to measure global coagulation kinetics, to ensure specific coagulation abnormalities are identified and treated in real-time.

### Machine Learning Methods Indicate Key Targets For Treatment

By integrating patient test results with machine learning algorithms, we were able to quantify key clinical and biomedical parameters which predict mortality and may be indicative of therapeutic targets. These algorithms identified D-dimer, PTT, clotting rate, fibrinogen concentration, and head abbreviated injury score (AIS) as the 5 most important markers predicting mortality in our trauma patient cohort. It is important to note that although PT and PTT (assays commonly used in clinical practice) broadly reflect hemostatic function, there is low correlation with clotting rate (−0.19 and -0.23 respectively) which is of great clinical significance because clotting rate is directly correlated with blood loss, development of shock, acidosis and ultimately death **(Figure S3)**. This further highlights the precision and value of viscoelastic assays in measuring clinically meaningful parameters. These predictive assessments suggest potential therapeutic targets such as immediate antifibrinolytic therapy administration in the pre-hospital phase of care, and early fibrinogen replacement informed by consumption rates to ensure clot formation and stability.

Our results are subject to several limitations. This study design only allows for a snapshot assessment of the coagulation abnormalities present following traumatic injury. Though patients had blood drawn upon admission to the resuscitation area, time of blood draw from time of injury varied from patient to patient. Due to this, the rate at which coagulation abnormalities occur and the point across the injury timeline varies between patients results in some heterogeneity which could not be accounted for with the current study design. Secondly, due to the strict inclusion/exclusion criteria established for this study, several potential subjects with life threatening injuries could not be enrolled due to all resources being dedicated to attempted recovery from terminal hemorrhagic shock. Third, due to established trauma triage protocols which have recognized over and under-triage rates, some patients who might have been appropriate subjects were not enrolled because they did not arrive to the resuscitation bay with the highest trauma activation level. Lastly, due to the lack of thromboelastography and thromboelastometry at the Robert Wood Johnson University Hospital, we were unable to directly compare our viscoelastic results with these assays. These limitations decreased the number of potential subjects and likely contributed to increased physiologic heterogeneity confounding our statistical analyses.

## CONCLUSIONS

Viscoelastic and biochemical assays play a key role in understanding the effect of injury severity on reducing clot stiffness, fibrinogen, and thrombin generation as well as increasing fibrinolysis. We believe that integrating hospital based viscoelastic assays to precisely assess coagulation system perturbations in real-time will enable targeted interventions that ultimately reduce post injury bleeding, transfusion requirements and mortality.

## Supporting information

VTutwiler_InjurySeverity_Supplemental

## Author Contributions

V. Tutwiler, A. Gosselin, C. Bargoud, J. Goswami, and J. Hanna designed experiments. A. Gosselin and M. Greenen performed experiments. V. Tutwiler, A. Gosselin, and M. Greenen analyzed data. C. Bargoud, S. Matthew, A. Toussaint, S. Coyle, and M. Macor obtained patient consent and assisted with patient sample delivery. A. Sawalkar and A. Krishnan performed and interpreted the machine learning algorithms and associated computational analyses. All authors contributed to the preparation of the manuscript.

## Acknowledgements

This work was supported by the National Institutes of Health (NIH) [NIH R00HL148646-01 (V.T.)], NIH [1T32GM135141-01A1 (A.G.)], New Jersey Commission For Spinal Cord Research [CSCR23IRG005 (V.T.)] and Rutgers University - New Brunswick, Office of the Vice Provost for Research, Research Ideation Seed Funding (V.T). A.K. was funded by US National Institutes of Health grants 1K08HG010061-01A1, 3UL1TR001085-04S1, and the MPN Research Foundation.

## Data Availability

Please contact the corresponding author.

