## Supplementary material for "Injury Severity is a Key Contributor to Coagulation Dysregulation and Fibrinogen Consumption": VTutwiler_InjurySeverity_Supplemental

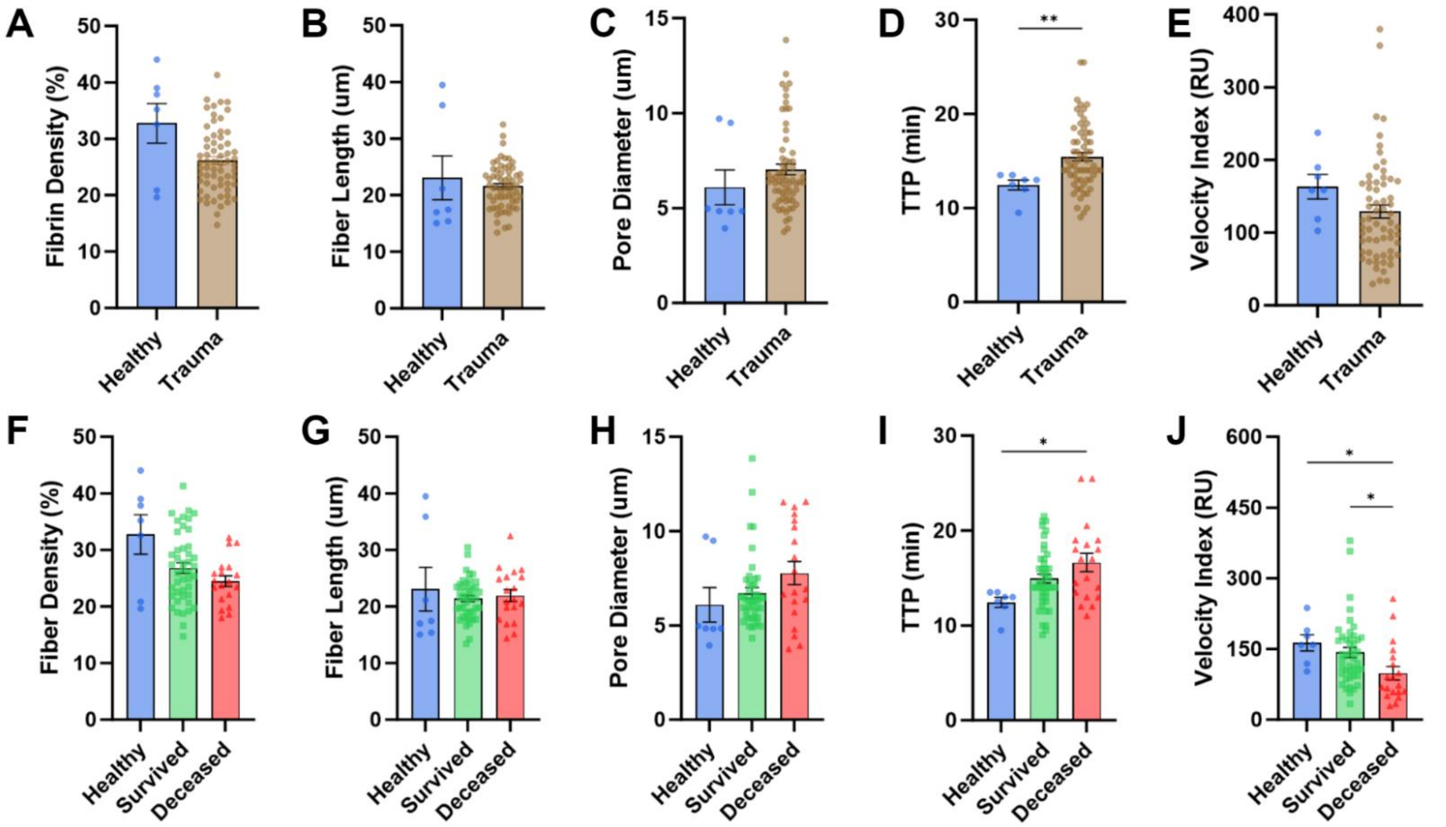

**<u>Figure S1: Trauma Patient and Healthy Donor Plasma Confocal and Thrombin Generation Properties</u>**
Comparison between confocal of healthy donors versus all trauma patients, as well as healthy donors versus survived and deceased patients. Including fibrin fiber density **(A,F)**, fiber length **(B,G)**, and pore diameter **(C,H)**. As well as thrombin generation parameters, time to peak thrombin **(D,I)** and velocity index **(E,J)**. *Significance between groups indicated by * $p < 0.05$, ** $< 0.01$.*

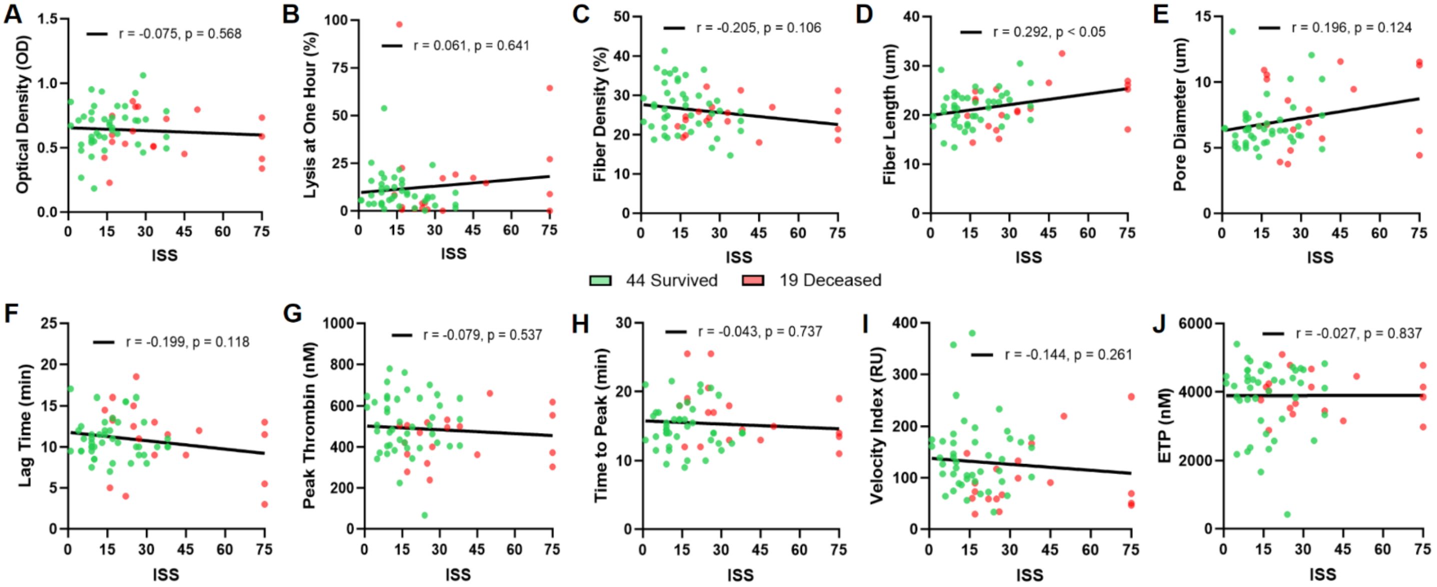

**Figure S2: Injury Severity Score Impact on Patient Coagulation Characteristics**

Correlation analysis of Injury Severity Score (ISS) and coagulation characteristics. Turbidity parameters, optical density **(A)**, and lysis at one hour **(B)**. Confocal analysis values, fibrin density **(C)**, fiber length **(D)** and pore diameter **(E)**. Thrombin generation parameters, lag time **(F)**, peak thrombin **(G)**, time to peak **(H)**, velocity index **(I)**, and endogenous thrombin potential **(J)**. *Significant correlations indicated by $p < 0.05$, r value indicates coefficient of correlation. Significance in t-test indicated by **** $p < 0.0001$.*

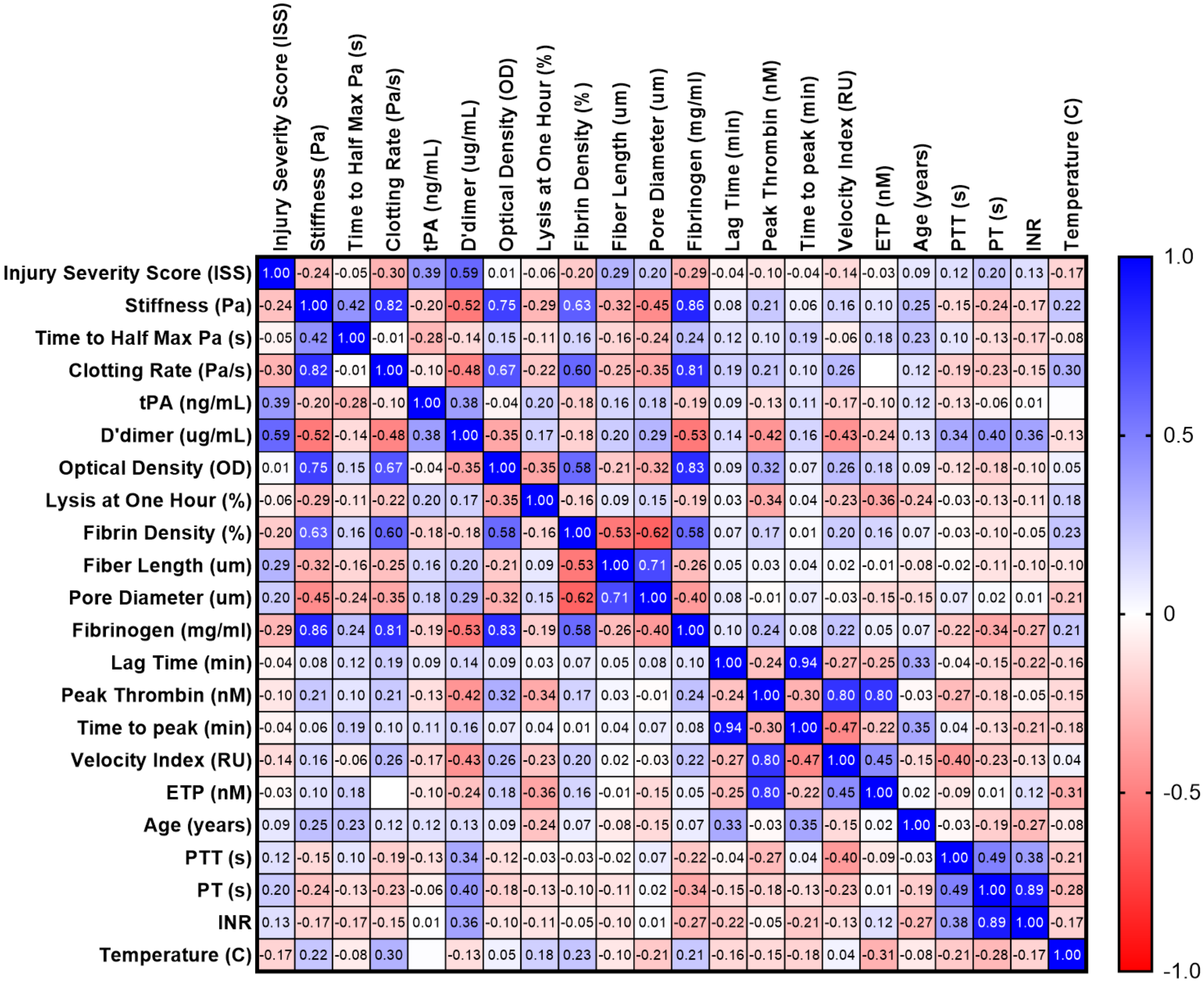

**Figure S3: Correlation Matrix Comparing Biomedical and Clinical Attributes**

Key Clinical and Biomedical variable correlation matrix indicating relationship between each characteristic. Dark blue indicates highly positive relationship, dark red indicates highly negative relationship, and white indicates no relationship.

**Table S1: Patient Inclusion and Exclusion Criteria**

| Inclusion Criteria | Exclusion Criteria |
|---|---|
| Patients > 18 years of age with any of the following<br>• Prehospital endotracheal intubation or assisted ventilation<br>• With respiratory rate < 10 or > 29 breaths per minute<br>Patients aged 18-64 years old with any of the following<br>• Systolic blood pressure < 90mmHg or sustained heart rate > 120 or < 40 beats per minute<br>Patients aged > 64 years old<br>• Systolic blood pressure < 100mmHg or sustained heart rate > 120 or < 40 beats per minute<br>Any patient with pre-hospital transfusion to maintain vital signs, with active hemorrhage, with glasgow coma score < 9, penetrating injury to the head, neck, torso, or extremities proximal to the elbow or knee or patients with crushed, amputated, degloved, mangled or pulseless extremities proximal to the wrist or ankle. | Patients from whom a sample could not be obtained prior to death.<br>Patients below 18 years of age.<br>Patients below 110 pounds.<br>Patients who are pregnant, have systemic anticoagulation, or antiplatelet medication at the time of injury.<br>Patients who are transferred from outside institutions or have a history of bleeding diathesis or hereditary coagulopathy. |

**Table S2:**

| **Clinical Parameters (85)** | **Biomedical Parameters (18)** |
|---|---|
| Age (years) | Stiffness (Pa) |
| Injury Mechanism | Time to Half Maximum Stiffness (s) |
| Injury Pattern | Clotting Rate (Pa/s) |
| Hemorrhage (yes/no) | Tissue Plasminogen Activator (pg/mL) |
| Traumatic Brain Injury (yes/no) | D-dimer (ng/mL) |
| Spinal Cord Injury (yes/no) | Optical Density (OD) |
| Blood Products (yes/no) | Time to Half Maximum Optical Density (s) |
| Blood Products (cumulative) | Clotting Rate (OD/s) |
| Sex | Lysis at One Hour (%) |
| Race | Fibrin Fiber Density (%) |
| Ethnicity | Fibrin Fiber Length (um) |
| Hypertension (yes/no) | Pore Diameter (um) |
| Chronic Kidney Disease (yes/no) | Fibrinogen (mg/mL) |
| Diabetes Mellitus (yes/no) | Lag Time (min) |
| Coronary Artery Disease (yes/no) | Peak Thrombin (nM) |
| Chronic Obstructive Pulmonary Disease (yes/no) | Time to Peak (min) |
| Bleeding Diathesis (yes/no) | Velocity Index |
| EtOH (mg/dL) | Endogenous Thrombin Potential (nM) |
| Hemoglobin (g/dL) | |
| Lactate (mmol/L) | |
| Utox Completed (yes/no) | |
| Utox Type 1 Amphetamines (yes/no) | |
| Utox Type 2 Barbiturates (yes/no) | |
| Utox Type 3 Benzodiazepine (yes/no) | |
| Utox Type 4 Cocaine (yes/no) | |
| Utox Type 5 Opiates (yes/no) | |
| Utox Type 6 PCP (yes/no) | |

| |
|---|
| Utox Type 7 THC (yes/no)<br>Utox Type 8 Oxycodone (yes/no)<br>Partial Thromboplastin Time (s)<br>Prothrombin Time (s)<br>International Normalized Ratio<br>D-dimer<br>Fibrinogen<br>Bicarbonate<br>Base<br>Tranexamic Acid (yes/no)<br>Pre-Hospital Blood (yes/no)<br>Fast (negative/positive)<br>Fast Loc 1 (RUQ)<br>Fast Loc 2 (LUQ)<br>Fast Loc 3 (Cardiac)<br>Fast Loc 4 (Pelvis)<br>Systolic Blood Pressure (mmHg)<br>Diastolic Blood Pressure (mmHg)<br>Heart Rate<br>Saturation (%)<br>Respiratory Rate<br>Temperature (C)<br>Glasgow Coma Score<br>Revised Trauma Score<br>Injury Severity Score (ISS)<br>TRISS<br>Abbreviated Injury Scale (AIS)<br>Cardiac Arrest (yes/no)<br>Blood within 24 hours (yes/no)<br>Cumulative BP<br>Packed Red Blood Cells<br>Fresh Frozen Plasma<br>Platelets<br>Cryoprecipitate<br>Massive Transfusion Protocol (yes/no)<br>Airway (needed intubation yes/no)<br>Airway type (endotracheal/surgical)<br>Chest tube (yes/no)<br>IR (yes/no)<br>OR (yes/no)<br>OR art (yes/no)<br>OR ven (yes/no)<br>Platelet count<br>TBI/SCI 1<br>TBI/SCI 2<br>AIS Head<br>AIS Face<br>AIS Neck<br>AIS Chest<br>AIS Abdomen<br>AIS Spine<br>AIS Upper<br>AIS Lower<br>AIS Extremities |

| |
|---|
| Deep Vein Thrombosis (yes/no)<br>Pulmonary Embolism (yes/no)<br>Stroke (yes/no)<br>Myocardial Infarction new (yes/no) |

**Table S3: Machine Learning Analysis Parameters**

| # | Method | Parameter | Accuracy (%) |
|---|---|---|---|
| 1 | Neural Net (NN) | Training Epochs: 100<br>Batch size: 16<br>Optimizer: Adam<br>Learning Rate: 0.005<br>Dropout: 50% at each layer<br>Activation Functions:<br>Hidden layers: ReLU<br>Output layer: Sigmoid<br>Architecture:<br>[103(Input) x 128 x 64 x 32 x 8 x 1 (Output)] | 79.62 |
| 2 | Support Vector Classifier (SVC) | Regularization parameter: 1.0<br>Linear Kernel, Shrinking heuristic enabled | 81 |
| 3 | Logistic Regression (LR) | Ridge Regularization used | 85 |
| 4 | Random Forest (RF) | Number of trees in the forest: 100<br>Number of features at each split: 4 | 85 |
| 5 | Random Forest (RF) – Hyper-tuned | Number of trees in the forest: 200<br>Number of parallel jobs: 10<br>Number of features to consider at each split: 20<br>Bootstrapped samples used<br>Out of bag samples used | 88 |

**Table S4: Healthy Donor and Trauma Patient Coagulation Average Characteristics**

| | Healthy | Trauma | Survived | Deceased |
|---|---|---|---|---|
| **Stiffness (Pa)** | 183.6 | 80.1 | 89.1 | 59.2 |
| **Clotting Rate (Pa/s)** | 0.24 | 0.19 | 0.23 | 0.12 |
| **Fibrinogen (mg/mL)** | 3.77 | 2.54 | 2.76 | 2.04 |
| **Optical Density (OD)** | 0.83 | 0.64 | 0.66 | 0.59 |
| **Lysis at One Hour (%)** | 3.46 | 12.12 | 10.2 | 16.38 |
| **D'dimer (ug/mL)** | 0.48 | 35.53 | 20.57 | 70.17 |
| **tPA (ng/mL)** | 2.38 | 13.01 | 9.98 | 20.01 |
| **Fiber Density (%)** | 32.74 | 26.11 | 26.83 | 24.46 |
| **Fiber Length (um)** | 23.06 | 21.56 | 21.4 | 21.95 |
| **Pore Diameter (um)** | 6.08 | 7.03 | 6.72 | 7.77 |
| **Lag Time (min)** | 8.43 | 11 | 10.97 | 11.09 |
| **Peak Thrombin (nM)** | 615 | 488 | 508 | 443 |
| **Time to Peak (min)** | 12.43 | 15.44 | 14.93 | 16.63 |
| **Velocity Index (RU)** | 163 | 129 | 142 | 98.2 |
| **ETP (nM)** | 4752 | 3891 | 3864 | 3954 |
